# SARS-CoV-2 Survival on Skin and its Transfer from Contaminated Surfaces

**DOI:** 10.1101/2024.09.18.613660

**Authors:** Ana K. Pitol, Samiksha Venkatesan, Siobhan Richards, Michael Hoptroff, Amitabha Majumdar, Grant Hughes

## Abstract

Understanding the transmission dynamics of SARS-CoV-2, particularly its transfer from contaminated surfaces (fomites) to human skin, is crucial for mitigating the spread of COVID-19. While extensive research has examined the persistence of SARS-CoV-2 on various surfaces, there is limited understanding of how efficiently it transfers to human skin, and how long it survives on the skin. This study investigates two key aspects of SARS-CoV-2 transmission: (1) the transfer efficiency of SARS-CoV-2 from non-porous (plastic and metal) and porous (cardboard) surfaces to a 3D human skin model (LabSkin), and (2) the survival of SARS-CoV-2 on the skin under different temperature conditions. First, we validated LabSkin as a suitable surrogate for human skin by comparing the transfer efficiency of the bacteriophage Phi 6 from surfaces to LabSkin and to human volunteers’ fingers. No significant differences were observed, confirming LabSkin’s suitability for these studies. Subsequently, the transfer of SARS-CoV-2 from surfaces to LabSkin was assessed, showing that plastic and metal surfaces had similar transfer efficiencies (∼13%), while no transfer occurred from cardboard once the inoculum had dried on the surface. Finally, the survival of SARS-CoV-2 on skin was assessed, showing a rapid decay at higher temperatures, with a half-life ranging from 2.8 to 17.8 hours depending on the temperature. These findings enhance our understanding of viral transmission via fomites and inform public health strategies to reduce the risk of SARS-CoV-2 transmission through surface contact.

## INTRODUCTION

Understanding the transmission pathways of infectious viruses is crucial for mitigating the spread of diseases (1). SARS-CoV-2, the virus responsible for COVID-19, is primarily transmitted through respiratory droplets and aerosols, but indirect contact through contaminated surfaces (fomites) plays a potentially significant role in its spread (2–4). While extensive research has been conducted on the persistence of SARS-CoV-2 on various surfaces, there is limited understanding of how efficiently it transfers from surfaces to human skin, particularly hands, and the subsequent risk of self-inoculation via hand-to-mouth contact (5).

Several studies have investigated the survival of SARS-CoV-2 on a range of surfaces, including plastic, metal, paper, and fabrics (6–12). The virus has been shown to remain viable for hours to days depending on the surface type, environmental conditions such as temperature and humidity, initial inoculum concentration and inoculum matrix (6–10,13–20). These findings have contributed significantly to our understanding of the virus’s environmental stability and the potential role of surfaces in transmission. However, while surface survival data is abundant, there is a noticeable gap in the literature when it comes to virus transfer dynamics—specifically, how efficiently SARS-CoV-2 transfers from contaminated surfaces to human skin and how long does SARS-CoV-2 survive on the skin (17,21,22). These are critical aspects of transmission, as the virus on the surfaces must transfer to the skin, typically the hands, before posing a risk of self-inoculation via mucous membranes (E.g., mouth, nose).

Virus transfer data is typically gathered from experiments involving volunteers, where bacteriophages are used as surrogates for pathogenic viruses (23–31). Bacteriophage Phi 6 is frequently employed as a surrogate for SARS-CoV-2 and other enveloped viruses in studies evaluating virus survival and transfer (32–36), due to its safety, low cost, and the fact that it does not require handling in high-containment facilities (37–39). However, research on the suitability of bacteriophages as surrogates for pathogenic viruses in studies evaluating virus transfer and survival has shown that bacteriophages can behave significantly differently from the viruses of concern (36,40). For instance, one study comparing the persistence of Phi 6 with SARS-CoV-2 on surfaces found that Phi 6 survived from 18 to 168 times longer than SARS-CoV-2 under the same experimental conditions (36). Therefore, it is crucial to directly assess the persistence and transfer of viruses using the specific virus of interest.

As an alternative to experiments conducted with volunteers and bacteriophages, various human skin substitutes have been proposed to study virus transfer and survival on the skin. These substitutes include human skin collected from surgeries (41,42), cadaver hands and arms (40), animal skin (17,43), and human skin models made with synthetic polymers (21,22,44). Three studies used the artificial skin model, VITRO-SKIN, to quantify the transfer of SARS-CoV-2 from objects to skin and its survival on the skin (21,22,44), while another two used pig skin as human skin model (17,43). While the aforementioned publications provide information on the transfer of SARS-CoV-2 from surfaces to skin models, they have a number of limitations. A study comparing virus transfer from environmental reservoirs to human skin showed that VITRO-SKIN is not a good skin surrogate for virus transfer studies, as it behaved significantly different than human skin (40). Additionally, pig skin and other animal models have not been validated as surrogates for human skin in virus transfer studies. Therefore, it is essential to evaluate human skin models that can serve as alternatives to human volunteers in studies aimed at assessing the transfer of pathogenic viruses to the skin.

Here, we investigate two key aspects of SARS-CoV-2 transmission dynamics: (1) the transfer of SARS-CoV-2 from surfaces to the skin, and (2) its survival on the skin. First, we assessed the suitability of using a 3D human skin model, LabSkin, to estimate virus transfer from surfaces to skin by comparing the transfer of a SARS-CoV-2 surrogate, bacteriophage Phi 6, from surfaces to the hands of human volunteers with its transfer from surfaces to LabSkin. After validating the human skin model, we quantified the transfer efficiency of SARS-CoV-2 from two non-porous surfaces—plastic and metal—and one porous surface, cardboard, to the human skin model. Finally, we quantified the persistence of SARS-CoV-2 on the skin. Through this research, we aim to fill a critical gap in understanding how viral transmission occurs via fomite contact, as well as how long SARS-CoV-2 persists on human skin. Our findings will not only enhance our understanding of viral transfer and persistence but also provide data necessary to develop evidence-based public health guidelines and design risk reduction strategies.

## METHODS

### SARS-CoV-2 production and quantification

Vero E6 cells (African green monkey kidney cells, ATCC) obtained from Public Health England were maintained at 37°C and 5% CO_2_ in Dulbecco’s Minimal Essential Medium (DMEM, Corning) with 10% fetal bovine serum (FBS, Sigma Aldrich) and 0.05 mg/ml gentamicin (Gibco). SARS-CoV-2 delta variant (SARS-CoV-2/human/GBR/Liv_273/2021, GenBank accession number OK392641) was maintained in Vero E6 cells. The number of infectious viral particles was quantified by standard plaque assay as previously described (36,45). Briefly, samples were serially diluted 10-fold and inoculated into the wells of a 24-well plate with Vero E6 cells at 80-100% confluence. One hour after infection, an agarose/growth media overlay (2% agarose in DMEM with 2% FBS) was applied to the cell monolayer. The plates were incubated for 72 hours at 37°C and 5% CO_2_. After incubation, the cells were fixed with 10% formalin (VWR International) and stained using a 0.25% solution of crystal violet (Sigma Aldrich). All SARS-CoV-2 work was conducted in a containment level 3 laboratory by personnel trained in the relevant codes of practice and standard operating procedures, who were vaccinated against SARS-CoV-2.

### Bacteriophage Phi 6 production and quantification

Bacteriophage Phi 6 (DSM 21518) and its host *Pseudomonas syringae* (DSM 21482) were obtained from the Deutsche Sammlung von Mikroorganismen und Zellkulturen (DSMZ). To culture the bacteriophage Phi 6, we used a protocol described elsewhere (46). Briefly, 100 mL of tryptic soy broth (TSB; Millipore) containing logarithmic phase *P. syringae* was inoculated with 100 μL of a 10^8^ PFU mL^-1^ stock of bacteriophage Phi 6 and incubated overnight. The next day, the media was centrifuged for 15 minutes at 5,000 rpm, and the supernatant was filtered using a 0.45-μm filter unit. Aliquots of the supernatant with a final concentration of 10^11^ PFU mL^-1^ were stored at 4°C for subsequent assays. To enumerate Phi 6, we used the standard double agar layer assay (46). Briefly, a 4 mL aliquot of soft TSB agar (0.5% Agar, Millipore) was inoculated with 100 μL of an overnight culture of *P. syringae* at a concentration of 10^9^ CFU mL^-1^ and 100 μL of the sample containing an unknown concentration of Phi6. Samples were mixed and poured onto hard agar (1.5% Agar) TSB plates and incubated at 25°C overnight. Negative controls (100 μL of an overnight culture of *P. syringae* with no bacteriophage) were included in each experiment. All dilutions were quantified in duplicates, and all experiments were performed in triplicates.

### LabSkin maintenance

Labskin 3D human constructs (LABSKIN) were maintained according to the manufacturer’s instructions. Briefly, the constructs were placed in a 12-well plate containing Labskin kit medium, with the upper part of the skin (stratum corneum) at the air interface and the bottom part (dermis) in contact with the culture medium. The Labskin constructs were incubated at 37°C and 5% CO_2_ for a maximum of 11 days before being used for experiments, with the culture medium replaced every second day.

### Surface preparation

The surfaces used for the transfer experiments were metal rods (stainless steel) alone or with one end covered by a circular piece (diameter = 10 mm) of the desired material (PVC plastic sheet, or cardboard). Metal rods and plastic surfaces were disinfected with 70% ethanol for 30 minutes. Cardboard surfaces were disinfected using UV light in the Class II Biological Safety Cabinet for 30 minutes.

### Ethical approval

The experimental design was approved by the Research Ethics Committee (REC) of the Liverpool School of Tropical Medicine (LSTM). All participants provided written informed consent after a thorough explanation of the study procedures, potential risks and benefits, and their right to withdraw at any time.

### Transfer of Phi 6 from surface to volunteers’ fingers and to artificial skin

To assess whether artificial skin (3D skin model, LabSkin) serves as a suitable surrogate for human skin in virus transfer experiments, we performed side-by-side comparisons using bacteriophage Phi 6 as a surrogate for pathogenic viruses. Bacteriophage Phi 6 is commonly used as a surrogate for enveloped viruses in various studies (33,36,47). We conducted six transfer events with 10 volunteers, totalling 60 transfers. These were compared to 60 transfers using LabSkin as substitute for the volunteers’ fingers. On the day of the experiments, humidity and temperature were measured before each experiment using the Testo 610 device. For the transfer events, steel rods were covered with plastic on one end. The plastic end was inoculated with 1 μL of bacteriophage Phi 6 at a concentration of 10^10^ PFU/mL. The inoculum was spread over the surface and allowed to dry for 1 hour. The rod was then placed on a balance, and a volunteer was asked to press one of their fingertips against the rod for 10 seconds, applying approximately 1.5 kPa of pressure (∼150 g/cm^2^). After each contact event, the surface of the rod and the skin were sampled by pipetting up and down 20 times over the contact area using cell culture media (DMEM, 2% FBS). This process was repeated six times, using the three middle fingers of both hands, with the thumbs serving as negative controls. The experiments were conducted across five days, with two volunteers per day.

A parallel comparison was conducted on the same days using LabSkin in place of volunteers’ fingers. The skin construct was placed on a balance, and the rod inoculated with Phi 6 was pressed against the skin for 10 seconds, applying 1.5 kPa of pressure. After the contact event, the rod and the skin were sampled similarly by pipetting up and down 20 times over the contact area using cell culture media. All samples were quantified on the same day using a standard plaque assay as previously described, with a method limit of quantification (LOQ) of 10 PFU.

Transfer efficiency was estimated using Equation 1, where *virus skin* (*PFU*) and *virus Surface* (*PFU*) are the number of viruses recovered from the skin and from the surface after the transfer event.

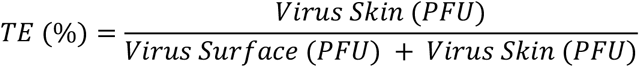

### Transfer of SARS-CoV-2 from surface to artificial skin

Having established that Labskin was an adequate surrogate for volunteers’ hands, we proceeded to evaluate the transfer efficiency of SARS-CoV-2 from surfaces of various materials to the skin. Three surface materials were evaluated: stainless steel, plastic, and cardboard. Similar to the experiment with bacteriophage Phi6, the rods (metal rods, or rods coated with plastic or cardboard) were inoculated with 1μL of SARS-CoV-2 at a concentration of 10^7^ PFU mL^-1^ and allowed to dry for a period of 1 hour. Before the contact event, the LabSkin construct was placed in a balance, to control the pressure applied. Contact between the rod and the skin was carried out by applying a pressure of ∼1,500 Pa of pressure (150 g/cm^2^) for 10 seconds. After the contact event, the surface and the skin were sampled by pipetting up and down 20 times the area of contact using cell culture media (DMEM, 2% FBS). Samples were quantified on the same day using standard plaque assay as previously described.

For the experiments of virus transfer from cardboard to skin, allowing the inoculum to dry for one hour on the cardboard resulted in a complete loss of the infective SARS-CoV-2. This meant that no virus was recovered from the surface or the skin before and after the transfer events. This can be explained by the adsorption of the virus to the permeable surface, a phenomenon previously observed on permeable materials such as cardboard (48). We therefore repeated the experiment with cardboard using a significantly smaller drying time (5 min), and a higher concentration of viruses (10^8^ PFU mL^-1^) to estimate virus transfer in a “worst case scenario”. Transfer efficiency of the virus was estimated using equation 1.

### SARS-CoV-2 survival on the skin

To evaluate SARS-CoV-2 survival on the skin, three droplets of SARS-CoV-2 in saliva at a concentration of 10^6^ PFU mL^-1^ were inoculated onto the Labskin construct. Saliva was used as the inoculation matrix due to its known influence on virus persistence on surfaces (16). The samples were allowed to dry in a microbiological cabinet, then placed inside a container and incubated at either 25°C or 37°C, or refrigerated at 4°C. Infectious virus was recovered from the skin over a time series by pipetting the skin up and down 20 times with 200 µL of DMEM supplemented with 2% FBS. Samples were frozen at -80°C until quantification. Viral quantification was performed using a standard plaque assay (36). Positive controls (viral stock) and negative controls (sampling LabSkin surface without viral inoculation) were included in the experiment. All experiments were conducted in triplicate.

### Data analysis

All statistical analyses were performed using R statistical software (version 4.3.3). A Student’s *t*-test was conducted to compare the transfer efficiency of bacteriophage Phi 6 from surfaces to volunteers’ fingers versus the transfer efficiency from surfaces to LabSkin. A Kruskal-Wallis test was employed to determine whether there was a statistically significant difference in the percent transfer efficiency of SARS-CoV-2 to the skin for the different surface materials tested (plastic, steel, cardboard), as the assumptions for parametric tests were not met. An ANCOVA test was used to predict the number of viruses on the skin as a function of time and temperature, and to determine whether there was a significant difference in the decay rate of SARS-CoV-2 as a function of temperature. Statistical significance for all the test was defined at an α value of <0.05. A linear model was utilized to estimate the half-life (*t*_1/2_) of SARS-CoV-2 on the skin for three different temperatures. Specifically, the concentration data of SARS-CoV-2 on the skin over time were log10-transformed. A linear regression model was then fitted to this transformed data. The slope of the resulting regression line, representing the decay rate, was used to calculate the half-life using the following equation:

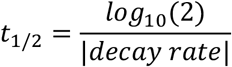

## RESULTS

### Transfer of Phi 6 from surface to volunteer’s fingers and to artificial skin

Transfer efficiency (TE, %) of bacteriophage Phi 6 from plastic surfaces to volunteers’ fingers was directly compared with transfers from surfaces to the 3D artificial skin, LabSkin. The temperature and humidity during the experiments were 20-21°C and 27-50%, respectively. All negative controls tested negative for the presence of bacteriophage Phi 6. Eighteen out of 120 data points were excluded from the analysis because the total number of viruses recovered (virus on the skin + virus on the surface) was below the limit of quantification. The mean transfer efficiency from plastic to skin was 12.2 ± 13.4 % (n=52) for LabSkin and 11.5 ± 12.1 % (n=50) for volunteers’ fingers. There was no statistically significant difference between the transfer efficiency of bacteriophage Phi 6 from surfaces to volunteers’ fingers compared to the transfer of virus from surfaces to LabSkin (Student’s *t*-test, *t*(100) = 0.25, p = 0.8), as observed in Figure 1.

**Figure 1.**
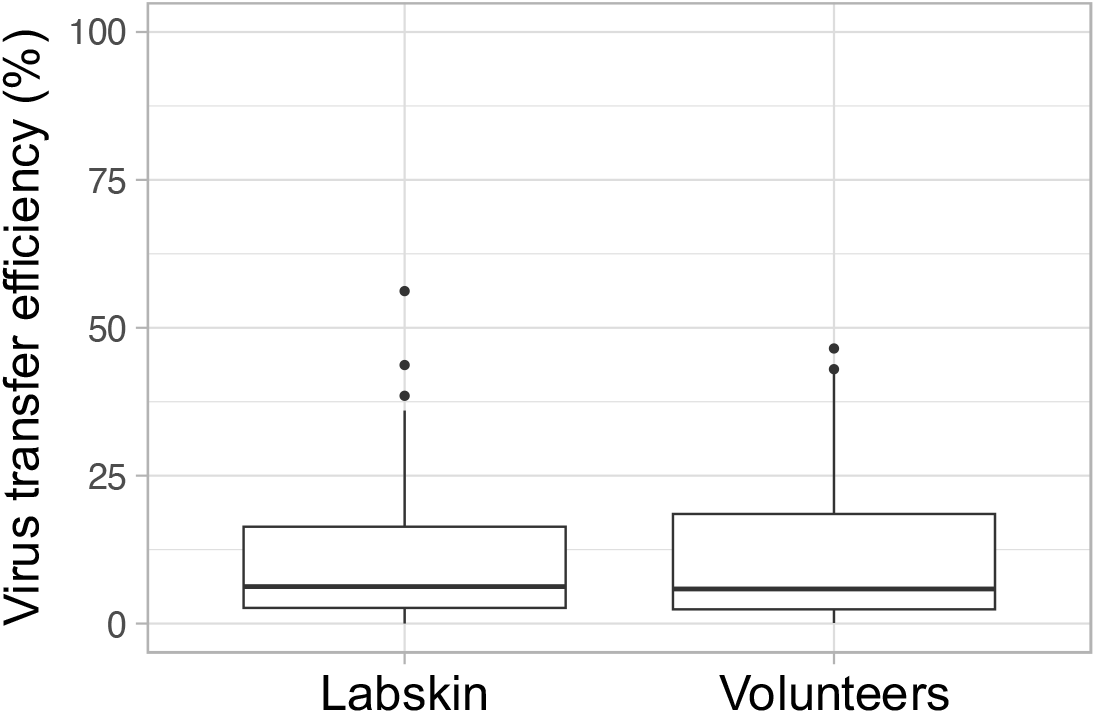
Transfer efficiency of bacteriophage Phi 6 from surface to skin as a function of skin type. The top and bottom of the box plots represent the 25th and 75th percentiles, the centerline represents the median value, and the whiskers extend to the highest and lowest concentrations. The total number of transfer events was 52 for Labskin and 50 for volunteers’ fingers (N=102).

### Transfer of SARS-CoV-2 from various surfaces to LabSkin

The temperature and humidity during the experiments ranged from 19-20°C and 26-30%, respectively. A total of 78 transfer events were conducted with SARS-CoV-2: 24 transfers from plastic surfaces, 24 from metal surfaces, and 30 from cardboard surfaces out of which 24 were wet transfers and 6 were dry transfers (Table 1). Nine out of the 78 data points were excluded from the analysis because the total number of viruses recovered (virus on the skin + virus on the surface) was below the limit of quantification (Table 1). The surface material significantly influenced virus transfer to the skin (Kruskal-Wallis, *H*(2) = 15.6, p<0.001). For the transfer of SARS-CoV-2 from cardboard to the skin, allowing the virus to dry on the cardboard for one hour resulted in no virus recovery from the surface. This indicates that the virus was completely adsorbed by the cardboard, rendering it unrecoverable after the one-hour period. The transfer efficiency (TE, %) for SARS-CoV-2 from plastic and metal surfaces to the skin was 13.7 ± 17.2 % and 13.6 ± 28.0 %, respectively (mean ± SD). The TE for cardboard was 0% when the virus was allowed to dry for 1 hour. Conversely, when the virus was left on the cardboard surface for only 5 minutes, a TE of 3.4 ± 10.5 % was observed (Table 1). While some SARS-CoV-2 transfer from wet cardboard to skin was detected shortly after inoculation (wet transfer), most of the transfer efficiencies from cardboard to skin were 0 even for wet transfers (16 out of 20 transfers).

**Table 1.** Transfer efficiency of SARS-CoV-2 from surfaces to skin.

| Surface | Inoculum |  | Transfer efficiency |  |  | Notes |
| --- | --- | --- | --- | --- | --- | --- |
|  | Concentration (PFU/mL) | Drying time (min) | n | Mean (SD) | Median (95% CI) |  |
| Plastic | $1 \times 10^7$ | 60 | 19 | 13.7 (17.2) | 7.9 (6.0, 21.5) | Dry transfer |
| Metal | $1 \times 10^7$ | 60 | 24 | 13.6 (28) | 1.6 (2.5, 24.8) | Dry transfer |
| Cardboard | $1 \times 10^7$ | 60 | 6 | NA | NA | Dry transfer |
| Cardboard | $1 \times 10^8$ | 5 | 20 | 3.4 (10.5) | 0 (-1.2, 8.0) | Wet transfer |

Note: For cardboard dry transfer we were unable to recover any virus from both, the surface and the skin, therefore no TE could be estimated (NA). A total of 24 transfer events were conducted for plastic, metal – dry transfer, and cardboard – wet transfer. Out of these, five data points for plastic and four data points for cardboard were excluded from the analysis, as the total number of virus recovered (virus on the surface + virus on the skin) fell below the limit of quantification.

The transfer efficiencies of SARS-CoV-2 from surfaces to skin in our study fell within the range reported in other studies for other viruses (Figure 2, SI Table 1). In Figure 2, we compare the results obtained from our experiments with those reported elsewhere, gathering information from similar experiments on virus transfer. The data included in Figure 2 was collected from literature data on similar studies conducted with volunteers and various viruses at similar temperature and humidity conditions, where “dry” transfer events were conducted (the inoculum was allowed to dry on the surface before the contact event). Significant variation can be observed between different studies as well as within the same studies, as indicated by the standard deviations of the reported data. The difference observed between studies may be attributed to various factors, which include variation in the methodology used to obtain the data as well as environmental conditions. More information on virus transfer data reported in Figure 2 can be found in the supporting information (SI Table 1).

**Figure 2.**
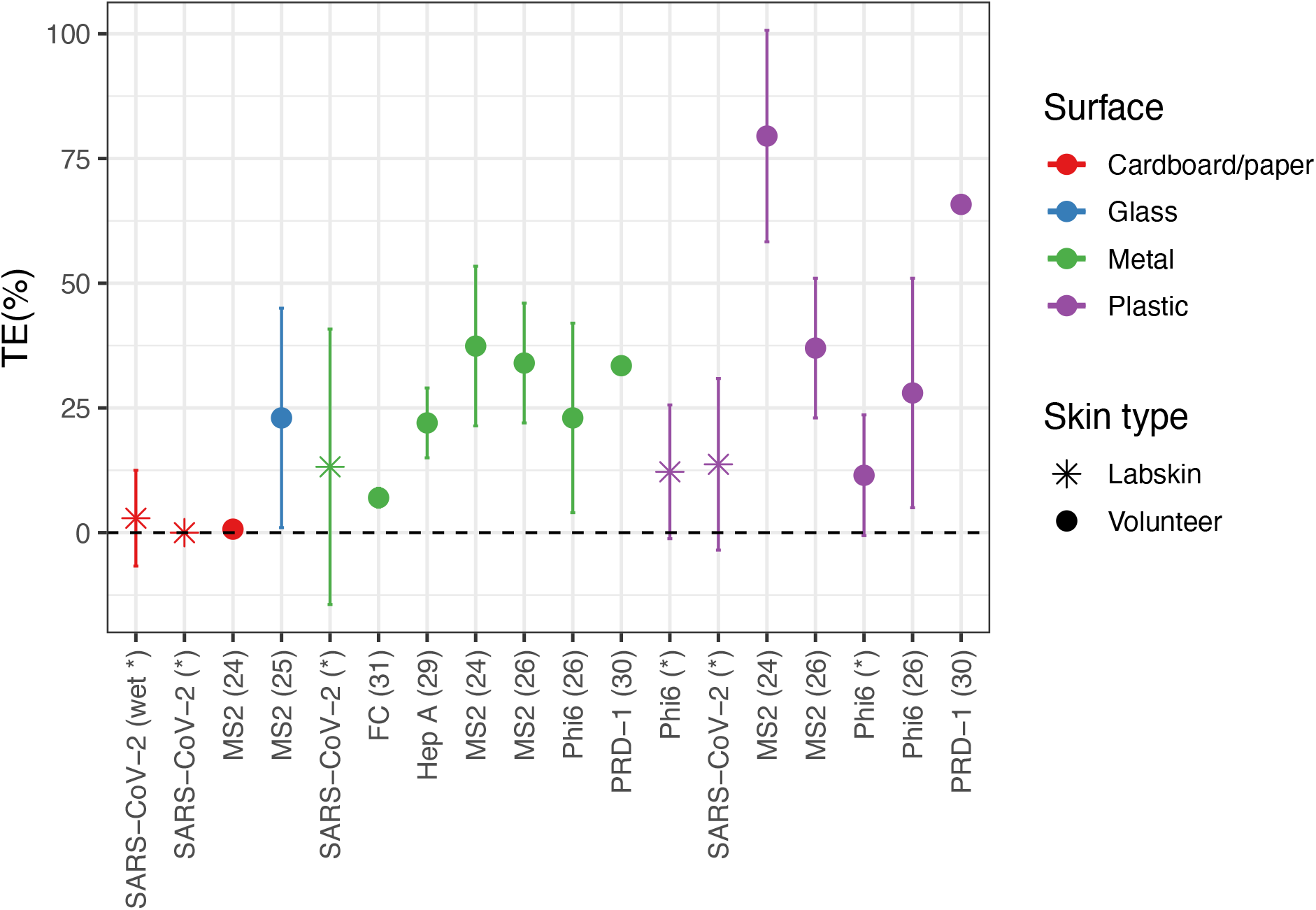
Transfer Efficiency of Viruses from Surfaces to Skin. Data points show the mean transfer efficiency (TE, %) of different viruses from metal (blue), plastic (purple), glass (green), and cardboard/paper (red) surfaces to skin. When available, the standard deviation of the data is shown. The skin type is represented by the shape: LabSkin is depicted as a star, and volunteers’ fingers as filled circles. The x-axis describes the virus used in the experiments, with the reference to the original manuscript in parentheses. Data from this study are shown with an asterisk instead of a reference number. All data reported in the plot, except in one case—specified by the word “wet” in parentheses—was collected from dry transfer events, where the surfaces were allowed to dry before the surface-to-skin contact. A detailed description of the studies reported in the figure can be found in the Supporting Information, Table S1.

### SARS-CoV-2 survival on artificial skin

The half-life of SARS-CoV-2 on the skin varied from 2.8 to 17.8 hours, for incubation temperatures between 4 and 37 °C. Temperature significantly influenced the survival of SARS-CoV-2 on the skin (ANCOVA, *F*(3,86)=15.9, p=0.03, *n*^*2*^= 0.72), indicating that the virus decay rate varied with different temperatures (Figure 3). The half-life of SARS-CoV-2 on the skin was 17.8 hours at 4°C, 4.1 hours at 25°C, and 2.8 hours at 37°C (Table 2).

**Table 2.**
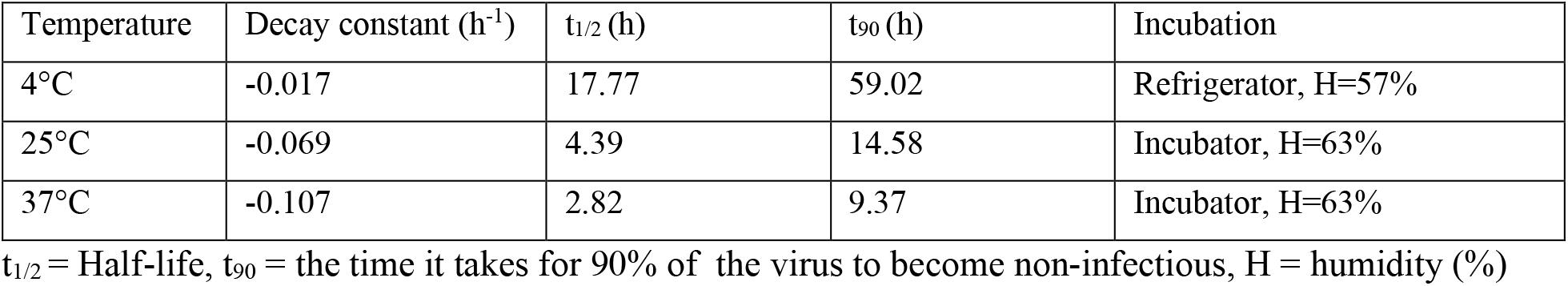
SARS-CoV-2 survival on the skin as a function of temperature.

| Temperature | Decay constant ($\text{h}^{-1}$) | $t_{1/2}$ (h) | $t_{90}$ (h) | Incubation |
| --- | --- | --- | --- | --- |
| 4°C | -0.017 | 17.77 | 59.02 | Refrigerator, H=57% |
| 25°C | -0.069 | 4.39 | 14.58 | Incubator, H=63% |
| 37°C | -0.107 | 2.82 | 9.37 | Incubator, H=63% |

$t_{1/2}$ = Half-life, $t_{90}$ = the time it takes for 90% of the virus to become non-infectious, H = humidity (%)

**Figure 3.**
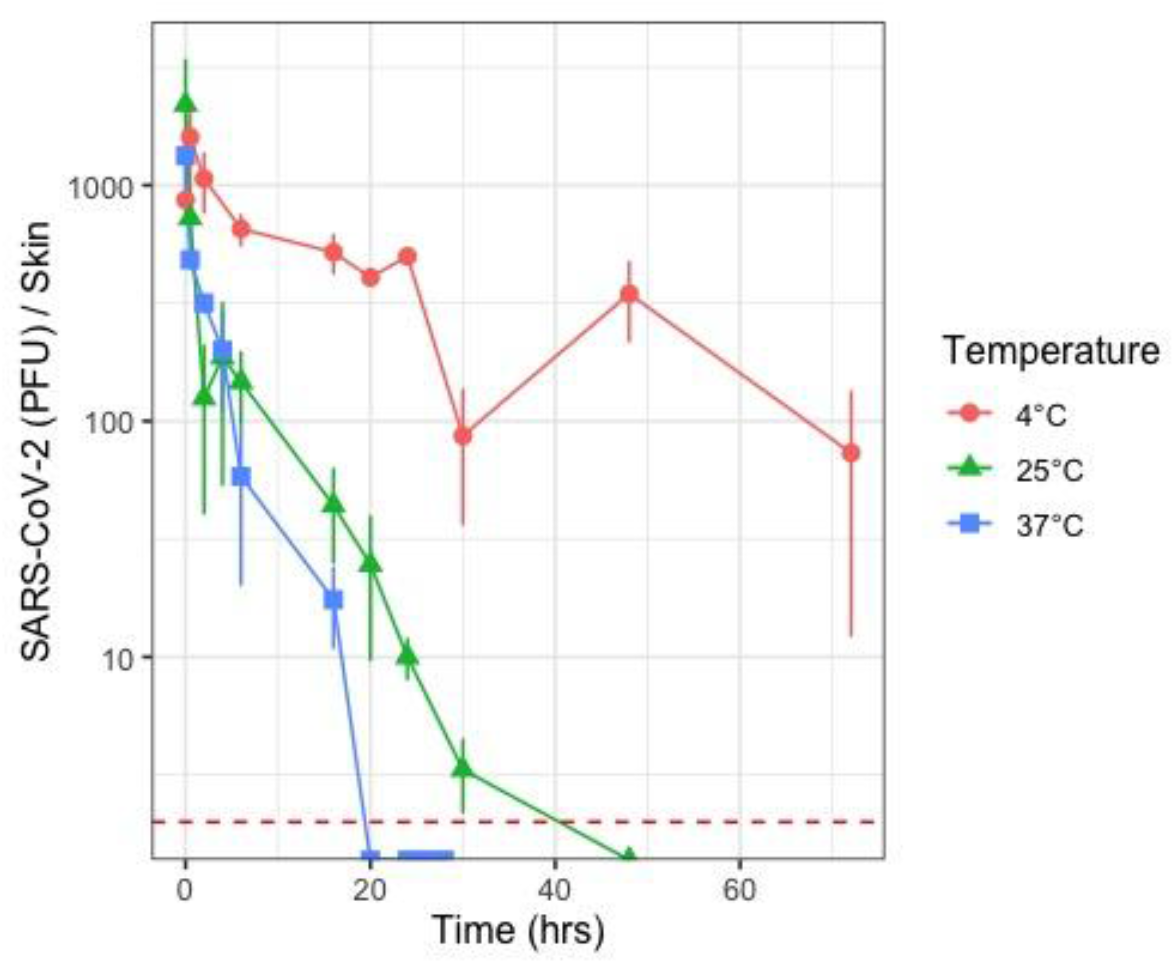
Survival of SARS-CoV-2 in skin as a function of temperature. Data shown is the average of three replicates ± standard deviation. Red dotted line represents the limit of detection of the assay.

## DISCUSSION

Virus transfer data is typically gathered from experiments involving volunteers, where bacteriophages are used as surrogates for pathogenic viruses (23–31). Alternatively, substitutes have been proposed to collect data on virus transfer and survival using the virus of interest in high-containment facilities, utilizing various human skin types that do not require the participation of volunteers. One study compared virus transfer to volunteers with virus transfer to cadaver hands and found no significant difference in transfer efficiencies (40). Alternatively, human skin obtained from surgeries is often used (41,42,49). Although the use of cadaver hands and human skin may yield results similar to those observed with volunteers, human skin and human hands are expensive, require ethical approval, and may lead to non-reproducible results. In this study, we conducted preliminary experiments comparing SARS-CoV-2 transfer from surfaces to two 3D human skin models (LabSkin and MatTek) and a synthetic skin model (VITRO-SKIN) with transfer to human skin obtained from surgeries (CTISkin). The results from the preliminary study indicated that LabSkin closely resembled human skin obtained from surgeries in virus transfer studies (SI, Figure 1). In addition to this, LabSkin model was thicker and therefore more robust for transfer experiments, which made it a more adequate alternative than the other skin models. Subsequently, we performed side-by-side comparisons of virus transfer from surfaces to volunteers’ fingers and to LabSkin, finding no significant difference in transfer efficiency between LabSkin and human hands. Therefore, LabSkin can be considered a suitable surrogate for human skin in transfer studies, offering a reproducible alternative to volunteers.

The transfer of SARS-CoV-2 from surfaces to LabSkin was significantly higher on non-porous surfaces (plastic, metal) compared to porous ones (cardboard). No virus was recovered from the cardboard after the virus was allowed to dry for one hour, so transfer efficiency could not be estimated for dry transfers. It has been reported that viruses decay rapidly on porous surfaces (14,15), and that the transfer efficiency of viruses from porous materials is significantly lower than from non-porous materials under various environmental conditions (17,22,24,30,31). Hosseini *et al*. highlighted that it is not the material’s porosity per se, but its permeability—or the ability of liquid to move across the material—that reduces virus infectivity on porous materials (48). They proposed three hypotheses to explain this phenomenon: 1) the material’s permeability increases the difficulty of recovering the virus from the surface, 2) it decreases the drying time, thereby accelerating viral decay, and 3) it increases the contact area between the virus and active ingredients on the surface, potentially inactivating the virus.

Similar findings of low SARS-CoV-2 recovery from porous materials have been reported in other studies. For instance, Johnson *et al*. (17) studied SARS-CoV-2 survival and transfer on cardboard using pig skin as human skin surrogate. In their study, they were able to recover only 11% of the inoculated SARS-CoV-2 from the cardboard three minutes after inoculation and could not recover any virus after 30 minutes (17). Pan *et al*. studied SARS-CoV-2 transfer from face masks to the artificial skin, VITRO-SKIN. They contaminated various facemasks (N95 respirator, surgical mask, polyester mask, cotton masks) with aerosols containing SARS-CoV-2 and subsequently performed transfer efficiency experiments by pressing the skin against the masks and quantifying virus transfer. They showed that no infectious virus was transferred from the masks to the skin (44). This is consistent with our data that shows that transfer efficiencies are low for permeable materials.

Our findings on SARS-CoV-2 transfer from surfaces to the skin align with previously reported data (Figure 2), though direct comparisons are challenging. The variability in transfer estimates across different studies can be attributed to differences in methodology (e.g., contact pressure, contact time), environmental conditions (e.g., temperature, humidity), the virus used, and the methods employed to calculate the transfer efficiencies. Transfer efficiencies reported in the literature have been calculated using three different approaches: 1) assuming the total virus count is the sum of the virus on the surface plus the virus on the skin (25,26,30), 2) using a control surface to estimate the total virus count (17,23,24,31), or 3) assuming that the total virus count corresponds to the initial inoculum deposited on the surface (21,22,27,50). The first and second methods are comparable, as they account for viruses lost during the drying process and those irreversibly adsorbed onto the surface. However, it is not feasible to compare these results with those obtained using the third method, where the total virus count is assumed to be equal to the initial inoculum (inoculation volume multiplied by inoculum concentration). This is because the transfer efficiency calculated in this manner is highly dependent on the time elapsed post-inoculation, the inoculum size and the drying conditions, incorporating information on virus transfer, virus decay, and irreversible adsorption. For instance, if 100 viruses are inoculated onto a surface, and after 30 minutes, 10 viruses are recovered from the skin following a transfer event, it remains unclear whether the transfer efficiency is 10% or if the actual transfer efficiency was higher, with the remaining viruses having decayed on the surface prior to the transfer event (e.g., if half of the viruses decayed, leaving 50, and 20% of those transferred to the skin). Therefore, this third methodology does not allow researchers to disentangle the different phenomena affecting the number of viruses recovered from the skin after contact. As a result, in this study, we excluded comparisons with other studies that calculated virus transfer using the initial inoculum as the total virus count.

In addition to assessing virus transfer, our study demonstrates that SARS-CoV-2 can remain infectious on human skin for hours to days, with its survival significantly influenced by temperature. Specifically, we observed that higher temperatures result in reduced viral half-lives, which aligns with existing literature indicating lower survival rates of SARS-CoV-2 at elevated temperatures (7,9,14,43). The impact of temperature on the persistence of viruses, including SARS-CoV-2, across various surfaces, food, skin, and other environmental reservoirs has been documented in numerous studies (51,52). These studies consistently show that higher temperatures accelerate the decay of infectious virus. Furthermore, temperature is just one of several environmental factors, including humidity and UV exposure, that can significantly alter virus stability and its transmission in different contexts (52,53).

Our study had a number of limitations. First, while we recorded environmental humidity during the virus persistence and transfer experiments, we did not actively control it. Humidity is a well-known factor that can significantly influence both virus persistence and transfer (24,54,55). Therefore, more experiments are needed to evaluate SARS-CoV-2 transfer from surfaces to skin and its persistence on the skin at various environmental conditions. Additionally, in the virus transfer experiments, we evaluated a single methodology, using a contact time of 10 seconds and a contact pressure of 1.5 kPa. However, the duration of contact, the applied pressure, and the friction between the finger and the surface are all variables that can influence the transfer of viruses (29,56,57). Moreover, the efficiency of virus transfer is highly dependent on the wetness inoculum on the surface at the time of contact (29,46,50). In real-world scenarios, most virus transfers are likely to occur through “dry transfers,” as the inoculation of surfaces typically involves very small droplets that dry out rapidly (e.g., these produced while talking, coughing, etc.). This drying process significantly alters the dynamics of virus transfer, potentially reducing the transfer efficiency compared to scenarios where the inoculum remains wet.

In conclusion, this study provides important insights into the mechanisms of SARS-CoV-2 transmission via surfaces and its subsequent survival on human skin. By validating the use of a 3D human skin model, LabSkin, as a surrogate for actual human skin, we were able to accurately quantify virus transfer from various surfaces and assess its survival on the skin over time. Our findings suggest that non-porous surfaces, such as plastic and metal, facilitate greater virus transfer compared to porous surfaces like cardboard, where the virus rapidly penetrates the material, making the recovery of infectious virus infeasible within minutes. Additionally, we found that SARS-CoV-2 can persist on human skin for significant periods, with higher temperatures accelerating viral decay. These findings underscore the complexity of virus transmission dynamics and emphasize the critical role of environmental factors in influencing virus survival. While our study advances the understanding of these processes, it also points to the necessity for further research into how other environmental variables, such as humidity and the state of the virus inoculum, and transfer parameters such as contact pressure and contact friction may affect virus transmission in everyday settings.

## ACKNOWLEDGMENT

This work was supported by grants awarded to A.K.P. by the National Institute for Health & Social Care Research Health Protection Research Unit (NIHR HPRU) in Emerging and Zoonotic Infections (NIHR200907) and the Pandemic Institute (TPIMPX01). The Pandemic Institute is formed of seven founding partners: The University of Liverpool, Liverpool School of Tropical Medicine, Liverpool John Moores University, Liverpool City Council, Liverpool City Region Combined Authority, Liverpool University Hospital Foundation Trust, and Knowledge Quarter Liverpool. This work was additionally supported by UKRI grants (20197 and 85336) awarded to G.L.H. G.L.H. was further supported by the BBSRC (BB/T001240/1, BB/X018024/1, BB/V011278/1, BB/X018024/1, and BB/W018446/1), the EPSRC (V043811/1), a Royal Society Wolfson Fellowship (RSWF\R1\180013), the NIHR (NIHR2000907), and the Bill and Melinda Gates Foundation (INV-048598). The views expressed are those of the author(s) and not necessarily those of the Pandemic Institute.

## CONFLICT OF INTEREST

M.H. and A.M. are Unilever employees.

## REFERENCES

1. WHO. Modes of transmission of virus causing COVID-19: implications for IPC precaution recommendations [Internet]. 2020 [cited 2024 Aug 20]. Available from: https://www.who.int/news-room/commentaries/detail/modes-of-transmission-of-virus-causing-covid-19-implications-for-ipc-precaution-recommendations

2. National Center for Immunization and Respiratory Diseases (NCIRD), Division of Viral Diseases. Scientific Brief: SARS-CoV-2 Transmission. CDC COVID-19 Science Briefs [Internet]. 2021 May 7 [cited 2022 Nov 18]; Available from: https://www.ncbi.nlm.nih.gov/books/NBK570442/

3. Centers for Disease Control and Prevention (CDC). COVID-19 Science Brief: SARS-CoV-2 and Surface (Fomite) Transmission for Indoor Community Environments [Internet]. 2021 Apr. Available from: https://www.cdc.gov/coronavirus/2019-ncov/more/science-and-research/surface-transmission.html

4. World Health Organization (WHO). Recommendation to Member States to improve hand hygiene practices widely to help prevent the transmission of the COVID-19 virus. Interim guidance. 2020.

5. Pitol AK, Julian TR. Community Transmission of SARS-CoV-2 by Surfaces: Risks and Risk Reduction Strategies. Environ Sci Technol Lett [Internet]. 2021 Jan;2020.11.20.20220749. Available from: http://medrxiv.org/content/early/2020/11/23/2020.11.20.20220749.abstract

6. van Doremalen N, Bushmaker T, Morris DH, Holbrook MG, Gamble A, Williamson BN, et al. Aerosol and Surface Stability of SARS-CoV-2 as Compared with SARS-CoV-1. New England Journal of Medicine [Internet]. 2020 Apr 16 [cited 2022 Nov 23];382(16):1564–7. Available from: https://www.nejm.org/doi/full/10.1056/nejmc2004973

7. Matson MJ, Yinda CK, Seifert SN, Bushmaker T, Fischer RJ, Doremalen N Van, et al. Effect of Environmental Conditions on SARS-CoV-2 Stability in Human Nasal Mucus and Sputum. Emerg Infect Dis [Internet]. 2020 Sep 1 [cited 2022 Nov 23];26(9):2276. Available from: /pmc/articles/PMC7454058/

8. Liu Y, Li T, Deng Y, Liu S, Zhang D, Li H, et al. Stability of SARS-CoV-2 on environmental surfaces and in human excreta. J Hosp Infect [Internet]. 2021 Jan 1 [cited 2022 Nov 23];107:105. Available from: /pmc/articles/PMC7603996/

9. Kwon T, Gaudreault NN, Richt JA, Vilibic-Cavlek T, Savic V. Environmental Stability of SARS-CoV-2 on Different Types of Surfaces under Indoor and Seasonal Climate Conditions. Pathogens 2021, Vol 10, Page 227 [Internet]. 2021 Feb 18 [cited 2022 Nov 23];10(2):227. Available from: https://www.mdpi.com/2076-0817/10/2/227/htm

10. Pottage T, Garratt I, Onianwa O, Spencer A, Paton S, Verlander NQ, et al. A comparison of persistence of SARS-CoV-2 variants on stainless steel. Journal of Hospital Infection. 2021 Aug 1;114:163–6.

11. Persistence of Severe Acute Respiratory Syndrome Coronavirus 2 (SARS-CoV-2) Virus and Viral RNA in Relation to Surface Type and Contamination Concentration [Internet]. [cited 2023 Jul 31]. Available from: 10.1128/AEM.00526-21

12. Issues I of M (US) C on PPE for HP to PT of PI and OVRICR, Larson EL, Liverman CT. Preventing Transmission of Pandemic Influenza and Other Viral Respiratory Diseases. Preventing Transmission of Pandemic Influenza and Other Viral Respiratory Diseases: Personal Protective Equipment for Healthcare Personnel: Update 2010 [Internet]. 2011 May 26 [cited 2023 Jul 31];p1– 187. Available from: https://www.ncbi.nlm.nih.gov/books/NBK209584/

13. Kramer A, Schwebke I, Kampf G. How long do nosocomial pathogens persist on inanimate surfaces? A systematic review. BMC Infect Dis [Internet]. 2006 Aug 16 [cited 2023 Feb 2];6(1):1– 8. Available from: 10.1186/1471-2334-6-130

14. Riddell S, Goldie S, Hill A, Eagles D, Drew TW. The effect of temperature on persistence of SARS-CoV-2 on common surfaces. Virol J [Internet]. 2020 Oct 7 [cited 2022 Nov 23];17(1):1–7. Available from: 10.1186/s12985-020-01418-7

15. Chin AWH, Chu JTS, Perera MRA, Hui KPY, Yen HL, Chan MCW, et al. Stability of SARS-CoV-2 in different environmental conditions. Lancet Microbe [Internet]. 2020 May 1 [cited 2023 Jul 31];1(1):e10. Available from: /pmc/articles/PMC7214863/

16. Ratnesar-Shumate S, Williams G, Green B, Krause M, Holland B, Wood S, et al. Simulated Sunlight Rapidly Inactivates SARS-CoV-2 on Surfaces. J Infect Dis [Internet]. 2020 Jun 29 [cited 2022 Nov 20];222(2):214–22. Available from: https://academic.oup.com/jid/article/222/2/214/5841129

17. Johnson GT, Loehle C, Zhou SS, Chiossone C, Palumbo J, Wiegand P, et al. Evaluation of the Survivability of SARS-CoV-2 on Cardboard and Plastic Surfaces and the Transferability of Virus from Surface to Skin. Occupational Diseases and Environmental Medicine [Internet]. 2021 Mar 17 [cited 2022 Nov 23];9(2):63–73. Available from: http://www.scirp.org/journal/PaperInformation.aspx?PaperID=109355

18. Pastorino B, Touret F, Gilles M, de Lamballerie X, Charrel RN. Prolonged Infectivity of SARS-CoV-2 in Fomites. Emerg Infect Dis [Internet]. 2020 Sep 1 [cited 2022 Nov 20];26(9):2256. Available from: /pmc/articles/PMC7454106/

19. Kasloff SB, Leung A, Strong JE, Funk D, Cutts T. Stability of SARS-CoV-2 on critical personal protective equipment. Sci Rep [Internet]. 2021;11(1):1–7. Available from: 10.1038/s41598-020-80098-3

20. Paton S, Spencer A, Garratt I, Thompson KA, Dinesh I, Aranega-Bou P, et al. Persistence of severe acute respiratory syndrome coronavirus 2 (sars-cov-2) virus and viral rna in relation to surface type and contamination concentration. Appl Environ Microbiol [Internet]. 2021 Jun 1 [cited 2023 Jul 31];87(14):1–9. Available from: https://journals.asm.org/journal/aem

21. Todt D, Meister TL, Tamele B, Howes J, Paulmann D, Becker B, et al. A realistic transfer method reveals low risk of SARS-CoV-2 transmission via contaminated euro coins and banknotes. iScience. 2021 Aug 20;24(8):102908.

22. Behzadinasab S, Chin AW, Hosseini M, Poon LL, Ducker WA, Poon llmpoon L, et al. SARS-CoV-2 virus transfers to skin through contact with contaminated solids. medRxiv [Internet]. 2021; Available from: 10.1101/2021.04.24.21256044

23. Reed SE. An investigation of the possible transmission of Rhinovirus colds through indirect contact. J Hyg (Lond). 1975;75(2):249–58.

24. Lopez GU, Gerba CP, Tamimi AH, Kitajima M, Maxwell SL, Rose JB. Transfer efficiency of bacteria and viruses from porous and nonporous fomites to fingers under different relative humidity conditions. Appl Environ Microbiol. 2013;79:5728–34.

25. Julian TR, Leckie JO, Boehm AB. Virus transfer between fingerpads and fomites. J Appl Microbiol. 2010;109:1868–74.

26. Anderson CE, Boehm AB. Transfer Rate of Enveloped and Nonenveloped Viruses between Fingerpads and Surfaces. Appl Environ Microbiol [Internet]. 2021 Oct 1 [cited 2022 Nov 20];87(22). Available from: 10.1128/AEM.01215-21

27. Ansari S, Springthorpe V, Sattar S, Rivard S, Rahman M. Potential role of hands in the spread of respiratory viral infections: studies with human parainfluenza virus 3 and rhinovirus. J Clin Microbiol [Internet]. 1991;29(10):2115–9. Available from: http://jcm.asm.org/content/29/10/2115.full.pdf

28. Ansari SA, Sattar SA, Springthorpe VS, Wells GA, Tostowaryk W. In vivo protocol for testing efficacy of hand-washing agents against viruses and bacteria: Experiments with rotavirus and Escherichia coli. Appl Environ Microbiol. 1989;55(12):3113–8.

29. Mbithi JN, Springthorpe VS, Boulet JR, Sattar S a. Survival of hepatitis A virus on human hands and its transfer on contact with animate and inanimate surfaces. J Clin Microbiol. 1992;30:757–63.

30. Rusin P, Maxwell S, Gerba C. Comparative surface-to-hand and fingertip-to-mouth transfer efficiency of gram-positive bacteria, gram-negative bacteria, and phage. J Appl Microbiol. 2002;93:585–92.

31. Bidawid S, Malik N, Adegbunrin O, Sattar SA, Farber JM. Norovirus cross-contamination during food handling and interruption of virus transfer by hand antisepsis: experiments with feline calicivirus as a surrogate. J Food Prot. 2004;67(1):103–9.

32. String GM, White MR, Gute DM, Mühlberger E, Lantagne DS. Selection of a SARS-CoV-2 Surrogate for Use in Surface Disinfection Efficacy Studies with Chlorine and Antimicrobial Surfaces. Environ Sci Technol Lett [Internet]. 2021 [cited 2022 Nov 20];8:995–1001. Available from: 10.1021/acs.estlett.1c00593

33. Baker CA, Gutierrez A, Gibson KE. Factors Impacting Persistence of Phi6 Bacteriophage, an Enveloped Virus Surrogate, on Fomite Surfaces. Appl Environ Microbiol [Internet]. 2022 Apr 1 [cited 2022 Nov 20];88(7). Available from: 10.1128/aem.02552-21

34. Baker CA, Gibson KE. Phi 6 recovery from inoculated fingerpads based on elution buffer and methodology. J Virol Methods. 2022 Jan 1;299:114307.

35. Fedorenko A, Grinberg M, Orevi T, Kashtan N. Survival of the enveloped bacteriophage Phi6 (a surrogate for SARS-CoV-2) in evaporated saliva microdroplets deposited on glass surfaces. Scientific Reports 2020 10:1 [Internet]. 2020 Dec 29 [cited 2022 Nov 20];10(1):1–10. Available from: https://www.nature.com/articles/s41598-020-79625-z

36. Pitol AK, Venkatesan S, Hoptroff M, Hughes GL. Persistence of SARS-CoV-2 and its surrogate, bacteriophage Phi6, on surfaces and in water. Appl Environ Microbiol [Internet]. 2023 Nov 29 [cited 2024 Jun 5];89(11). Available from: 10.1128/aem.01219-23

37. Gallandat K, Lantagne D. Selection of a Biosafety Level 1 (BSL-1) surrogate to evaluate surface disinfection efficacy in Ebola outbreaks: Comparison of four bacteriophages. PLoS One [Internet]. 2017 May 1 [cited 2022 Nov 20];12(5):e0177943. Available from: 10.1371/journal.pone.0177943

38. Aranha-Creado H, Brandwein H. Application of Bacteriophages as Surrogates for Mammalian Viruses: A Case for Use in Filter Validation Based on Precedents and Current Practices in Medical and Environmental Virology. PDA J Pharm Sci Technol. 1999;53(2).

39. Aquino De Carvalho N, Stachler EN, Cimabue N, Bibby K. Evaluation of Phi6 Persistence and Suitability as an Enveloped Virus Surrogate. Environ Sci Technol [Internet]. 2017 Aug 1 [cited 2022 Nov 20];51(15):8692–700. Available from: 10.1021/acs.est.7b01296

40. Pitol AK, Bischel HN, Boehm AB, Kohn T, Julian TR. Transfer of Enteric Viruses Adenovirus and Coxsackievirus and Bacteriophage MS2 from Liquid to Human Skin. Appl Environ Microbiol [Internet]. 2018;84(22):1–13. Available from: 10.1128/AEM.01809-18

41. Hirose R, Bandou R, Ikegaya H, Watanabe N, Yoshida T, Daidoji T, et al. Disinfectant effectiveness against SARS-CoV-2 and influenza viruses present on human skin: model-based evaluation. Clinical Microbiology and Infection. 2021 Jul 1;27(7):1042.e1-1042.e4.

42. Hirose R, Ikegaya H, Naito Y, Watanabe N, Yoshida T, Bandou R, et al. Survival of Severe Acute Respiratory Syndrome Coronavirus 2 (SARS-CoV-2) and Influenza Virus on Human Skin: Importance of Hand Hygiene in Coronavirus Disease 2019 (COVID-19). Clinical Infectious Diseases. 2021 Dec 6;73(11):e4329–35.

43. Harbourt DE, Haddow AD, Piper AE, Bloomfield H, Kearney BJ, Fetterer D, et al. Modeling the stability of severe acute respiratory syndrome coronavirus 2 (SARS-CoV-2) on skin, currency, and clothing. PLoS Negl Trop Dis [Internet]. 2020 Nov 1 [cited 2022 Nov 18];14(11):e0008831. Available from: 10.1371/journal.pntd.0008831

44. Pan J, Gmati S, Roper BA, Prussin AJ, Hawks SA, Whittington AR, et al. Stability of Aerosolized SARS-CoV-2 on Masks and Transfer to Skin. Environ Sci Technol [Internet]. 2023 Jul 18 [cited 2024 Jul 31];57(28):10193–200. Available from: 10.1021/acs.est.3c01581

45. Anderson ER, Hughes GL, Patterson EI. Inactivation of SARS-CoV-2 on surfaces and in solution with Virusend (TX-10), a novel disinfectant. Access Microbiol. 2021;3(4):10–3.

46. Pitol AK, Bischel HN, Kohn T, Julian TR. Virus Transfer at the Skin-Liquid Interface. Environ Sci Technol. 2017 Dec 19;51(24):14417–25.

47. Ye Y, Ellenberg RM, Graham KE, Wigginton KR. Survivability, Partitioning, and Recovery of Enveloped Viruses in Untreated Municipal Wastewater. Environ Sci Technol [Internet]. 2016 May 17 [cited 2022 Nov 20];50(10):5077–85. Available from: 10.1021/acs.est.6b00876

48. Hosseini M, Poon LLM, Chin AWH, Ducker WA. Effect of Surface Porosity on SARS-CoV-2 Fomite Infectivity. ACS Omega [Internet]. 2022 Jun 7 [cited 2023 Jul 31];7(22):18238–46. Available from: 10.1021/acsomega.1c06880

49. Hirose R, Bandou R, Ikegaya H, Watanabe N, Yoshida T, Daidoji T, et al. Disinfectant effectiveness against SARS-CoV-2 and influenza viruses present on human skin: model-based evaluation. Clinical Microbiology and Infection. 2021 Jul 1;27(7):1042.e1-1042.e4.

50. Ansari S a., Sattar S a., Springthorpe VS, Wells G a., Tostowaryk W. Rotavirus survival on human hands and transfer of infectious virus to animate and nonporous inanimate surfaces. J Clin Microbiol. 1988;26:1513–8.

51. Boone SA, Gerba CP. Significance of Fomites in the Spread of Respiratory and Enteric Viral Disease. Appl Environ Microbiol [Internet]. 2007 [cited 2024 Aug 28];73(6):1687–96. Available from: https://journals.asm.org/journal/aem

52. Pirtle EC, Beran GW. Virus survival in the environment. Rev sci tech Off int Epiz. 1991;10(3):733– 48.

53. Tang JW. The effect of environmental parameters on the survival of airborne infectious agents. J R Soc Interface [Internet]. 2009 Dec 12 [cited 2024 Aug 28];6(Suppl 6):S737. Available from: /pmc/articles/PMC2843949/

54. Walker MD, Vincent JC, Benson L, Stone CA, Harris G, Ambler RE, et al. Effect of Relative Humidity on Transfer of Aerosol-Deposited Artificial and Human Saliva from Surfaces to Artificial Finger-Pads. Viruses [Internet]. 2022 May 1 [cited 2023 Jul 3];14(5):1048. Available from: https://www.mdpi.com/1999-4915/14/5/1048/htm

55. Walker MD, Vincent JC, Benson L, Stone CA, Harris G, Ambler RE, et al. Effect of Relative Humidity on Transfer of Aerosol-Deposited Artificial and Human Saliva from Surfaces to Artificial Finger-Pads. Viruses [Internet]. 2022 May 1 [cited 2024 Aug 28];14(5):1048. Available from: https://www.mdpi.com/1999-4915/14/5/1048/htm

56. Baker CA, Hamilton AN, Chandran S, Poncet AM, Gibson KE. Transfer of Phi6 bacteriophage between human skin and surfaces common to consumer-facing environments. J Appl Microbiol [Internet]. 2022 Dec 1 [cited 2024 Jul 31];133(6):3719–27. Available from: 10.1111/jam.15809

57. Zhao P, Chan PT, Gao Y, Lai HW, Zhang T, Li Y. Physical factors that affect microbial transfer during surface touch. Build Environ. 2019 Jul 1;158:28–38.

