## Supplemental Information for "SARS-CoV-2 Survival on Skin and its Transfer from Contaminated Surfaces"

We compared the transfer efficiency of SARS-CoV-2 from surface (metallic rod) to human skin (ex-vivo skin, CTISkin) and to various skin models. The models include two full thickness 3D human skin models: LabSkin (LABSKIN) and EpiDermFT (MatTek) and one synthetic skin surrogate, Vitro-Skin (VITRO-SKIN). All the skins were maintained according to the manufacturer's instructions.

To perform the transfer events, we used the protocol described in the main article. Briefly, we added a droplet of 2μL of SARS-CoV-2 solution, containing ~10^7^ PFU/mL, to the centre of a metallic rod and allowed the inoculum to dry. After drying time (1 hour), we placed the skin (VitroSkin, LabSkin, MatTek, or ex-vivo skin) on a balance placed inside the biological safety cabinet and followed with a 10 second contact event (150 ± 20 gr) between the metallic rod and the skin. After the transfer, we recovered the virus on from the surface and the skin by pipetting up and down 15 times using culture media (DMEM supplemented with 2% FBS). After finishing with all the skin transfers, we serial dilute the samples and plated them. Samples were quantified using standard plaque assays. Transfer efficiency was estimated using the following formula:

$$TE (\%)=\frac{Virus Skin (PFU)}{Virus Surface (PFU) + Virus Skin (PFU)}$$

where $Virus Skin (PFU)$ and $Virus Surface (PFU)$ are the number of viruses recovered from the skin and from the surface after the transfer event.

**Results**: SARS-CoV-2 transfer to ex-vivo human skin was similar to transfer to LabSkin but different to the transfer to MatTek and Vitro-Skin (Figure S1). Based on these data, we selected Labskin as a model for human skin in the proposed experiments given its similarity to human skin explants.

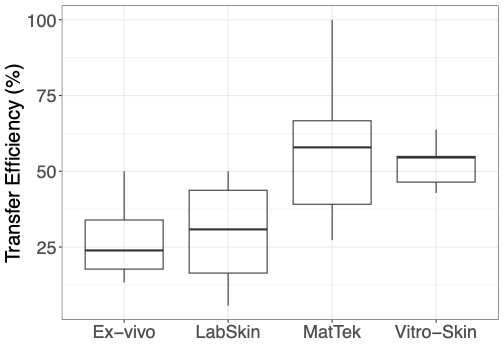

**Figure S1. Transfer Efficiency of SARS-CoV-2 from Surface to Skin as a Function of Skin Type.** The box plots show the distribution of transfer efficiency (TE), with the top and bottom edges representing the 25th and 75th percentiles, the centerline indicating the median value, and the whiskers extending to the highest and lowest observed values. The total number of transfer events was four for ex vivo skin and LabSkin, nine for MatTek, and five for VitroSkin. The sample size was determined by the availability of the different skin types. Experiments were conducted side-by-side to minimize variation in TE due to environmental factors such as humidity and temperature.

**Table S1. Virus Transfer Efficiency (TE, %) from Surface to Skin.**

This table presents the TE data that directly compares with the conditions used in this study, specifically dry transfer events, where the TE was estimated using a similar methodology than the one reported in the main manuscript.

| **Ref.** | **Virus** | **Surface** | **Skin** | **Drying**  **Time (min)** | **Contact time and contact pressure** | **Number of replicates** | **TE ± SD (%)** | **T and H** |
| --- | --- | --- | --- | --- | --- | --- | --- | --- |
| This study | Bacteriophage Phi 6 | Plastic | Volunteers’ fingers | 50 | 10 sec,  1,470 Pa | 50 | 11.5 ± 12.1 | 19-20 °C  26-30 % |
|  |  |  | Labskin | 52 |  | 52 | 12.2 ± 13.4 |  |
|  | SARS-CoV-2 | Plastic | Labskin | 24 |  | 24 | 13.7 ± 17.2 |  |
|  |  | Steel |  | 24 |  | 24 | 13.2 ± 27.6 |  |
|  |  | Cardboard* |  | 24 |  | 24 | 2.9 ± 9.6 |  |
| (1) | Bacteriophage MS2 | Steel | Volunteers’ fingers | 6 | 10 sec,  9,800 Pa | 6 | 37.4 ± 16 | 19-25 °C  40-65 % |
|  |  | Plastic |  | 6 |  | 6 | 79.5 ± 21.2 |  |
|  |  | Paper currency |  | 6 |  | 6 | 0.7 ± 0.5 |  |
| (2) | Bacteriophage MS2 | Glass | Volunteers’ fingers | 75 | 10 sec,  25,000 Pa | 75 | 25 ± 23 | 10-22 °C  40-65 % |
| (3) | Bacteriophage MS2 | Steel | Volunteers’ fingers | 30 | 10 sec,  2,450 Pa | 30 | 34 ± 12 | 21-22 °C  13-74 % |
|  |  | Plastic |  | 30 |  | 30 | 37 ± 14 |  |
|  | Bacteriophage Phi6 | Steel |  | 30 |  | 30 | 23 ± 19 |  |
|  |  | Plastic |  | 30 |  | 30 | 28 ± 23 |  |
| (4) | Bacteriophage PRD-1 | Steel | Volunteers’ fingers | NA | 10 sec,  NA | NA | 33.5 ± NA | NA |
|  |  | Plastic |  | NA |  | NA | 65.8 ± NA |  |
| (5) | Hepatitis A | Steel | Volunteers’ fingers | 6 | 10 sec,  9,800 and 1960 Pa | 6 | 22 ± 7 | 22 ± 2 °C  45 ± 5 % |
| (6) | Feline Calicivirus | Steel | Volunteers’ fingers | NA | 10 sec,  1960-3920 Pa | NA | 7 ± 1.9 | 24 ± 2 °C  45 % |

*Cardboard TE (%) reported here was evaluated at different drying time than plastic and steel.

T = Temperature, H = Humidity.

References:

1.	Lopez GU, Gerba CP, Tamimi AH, Kitajima M, Maxwell SL, Rose JB. Transfer efficiency of bacteria and viruses from porous and nonporous fomites to fingers under different relative humidity conditions. Appl Environ Microbiol. 2013;79:5728–34.

2.	Julian TR, Leckie JO, Boehm AB. Virus transfer between fingerpads and fomites. J Appl Microbiol. 2010;109:1868–74.

3.	Anderson CE, Boehm AB. Transfer Rate of Enveloped and Nonenveloped Viruses between Fingerpads and Surfaces. Appl Environ Microbiol [Internet]. 2021 Oct 1 [cited 2022 Nov 20];87(22). Available from: https://journals.asm.org/doi/10.1128/AEM.01215-21

4.	Rusin P, Maxwell S, Gerba C. Comparative surface-to-hand and fingertip-to-mouth transfer efficiency of gram-positive bacteria, gram-negative bacteria, and phage. J Appl Microbiol. 2002;93:585–92.

5.	Mbithi JN, Springthorpe VS, Boulet JR, Sattar S a. Survival of hepatitis A virus on human hands and its transfer on contact with animate and inanimate surfaces. J Clin Microbiol. 1992;30:757–63.

6.	Bidawid S, Malik N, Adegbunrin O, Sattar SA, Farber JM. Norovirus cross-contamination during food handling and interruption of virus transfer by hand antisepsis: experiments with feline calicivirus as a surrogate. J Food Prot. 2004;67(1):103–9.
